# Deep learning from harmonized peptide libraries enables retention time prediction of diverse post translational modifications

**DOI:** 10.1101/2023.05.30.542978

**Authors:** Damien Beau Wilburn, Ariana E. Shannon, Vic Spicer, Alicia L. Richards, Darien Yeung, Danielle L. Swaney, Oleg V. Krokhin, Brian C. Searle

## Abstract

In proteomics experiments, peptide retention time (RT) is an orthogonal property to fragmentation when assessing detection confidence. Advances in deep learning enable accurate RT prediction for any peptide from sequence alone, including those yet to be experimentally observed. Here we present Chronologer, an open-source software tool for rapid and accurate peptide RT prediction. Using new approaches to harmonize and false-discovery correct across independently collected datasets, Chronologer is built on a massive database with >2.2 million peptides including 10 common post-translational modification (PTM) types. By linking knowledge learned across diverse peptide chemistries, Chronologer predicts RTs with less than two-thirds the error of other deep learning tools. We show how RT for rare PTMs, such as OGlcNAc, can be learned with high accuracy using as few as 10-100 example peptides in newly harmonized datasets. This iteratively updatable workflow enables Chronologer to comprehensively predict RTs for PTM-marked peptides across entire proteomes.

## Introduction

Liquid chromatography (LC) is a common analytical technique that separates mixtures of compounds based on their physicochemical properties.^1^ In proteomic studies, LC is often used to separate proteins or peptides in complex biological mixtures, which simplifies analysis with mass spectrometry (MS) by reducing ionization competition and interference. Several separation chemistries exist for resolving analytes based on their charge (ion exchange), volume (size-exclusion), hydrophobicity/hydrophilicity, affinity to an immobilized ligand (e.g. antibody or metal ion), or in tandem for multidimensional separations (such as in techniques like MudPIT).^2^ In typical bottom-up proteomics experiments, the final peptide separation is performed directly inline with the mass spectrometer by reversed-phase (RP) chromatography where hydrophobicity is the principal driver of interactions with a stationary phase. When properly normalized, the retention time (RT) at which a peptide/protein elutes is an alternative measurement to mass spectra in assessing statistical confidence of peptide and protein detection. Accuracy in orthogonal measurements like RT is critical for confirming single peptide detections, which is necessary for measuring post-translational modification (PTM) containing peptides or human leukocyte antigens (HLAs).

Library search engines that incorporate RT scoring are common in data-independent acquisition (DIA) studies,^3–7^ and can improve searches in data-dependent acquisition (DDA) studies.^8^ Describing the relative hydrophobicity or elution time of a peptide has proved challenging due to the complicated physicochemical interactions that occur between C18 resins, background matrix, and peptides as they travel the length of the column. This is further confounded by experimental variation in instrument settings and column characteristics (packing, conditioning, etc.) that have observable influences on elution behavior. Hence, ideal libraries for searching MS data would include RTs measured using the same matrix, LC column, instrument, and buffers. The expense of experiment-specific RT libraries, which requires both greater instrument and time costs, has been mitigated by two main strategies: large-scale RT library re-use^9,10^ requiring alignment to map library RTs into experiment-specific space,^11,12^ and RT prediction directly from peptide sequences (reviewed in Krokhin 2012^13^).

Several RT prediction algorithms have been developed that range from simple linear regression models of bulk amino acid properties,^14^ to sophisticated biophysical models,^15^ and ultimately to deep learning methods.^16–20^ Deep learning is an extension of machine learning that uses large datasets to train highly parameter-rich models, where “deep” refers to both the number of hidden layers embedded within the neural networks and the scale of data necessary to train these large models.^21,22^ To predict RT, deep learning tools encode peptide sequences as a matrix of coefficients, then apply a series of learned matrix manipulations to derive a single elution time without any prior knowledge of chromatography. However, differences in columns, gradients, buffers, and LC systems can contribute to substantial deviations in RT for the same peptide. As such, current peptide RT prediction deep learning models have utilized one large dataset to train a single model. Newer approaches, such as DeepLC^19^ and AlphaPeptDeep,^20^ train multiple models based on different datasets and automatically select the model that best correlates with the user’s configuration. As an alternative, AlphaPeptDeep and AutoRT^18^ can use transfer learning to refine predictions derived from a small set of experiment-specific training data.

Machine learning assumes all training observations are “correct,” while proteomics data inherently has some degree of assumed error. Prosit^17^ solved this problem by training exclusively using a large library of synthetic human peptides as part of the ProteomeTools project.^23^ Unfortunately, large synthetic datasets are impossible to create for every experimental situation, which limits the accuracy of models produced from traditional false discovery rate (FDR)-controlled training datasets. The use of separate models for different data types produces other complicated ramifications. For example, AlphaPeptDeep has separate RT models for phosphorylated and unphosphorylated peptides that were trained on separate FDR-controlled datasets, which produce RTs configured for different experiments. This approach impedes the ability to predict how phosphorylated and unphosphorylated peptides elute in the same experiment. Building a new model for each training dataset reduces the impact of the most valuable aspect of deep learning: the ability to synthesize complex information in large-scale data. The performance of deep learning models is highly dependent on the breadth and scale of training data such that it would be preferable if one model were trained on a single, unified dataset with information on multiple modification types and peptide chemistries.

In this work, we present Chronologer, a new deep-learning architecture for learning RT. As part of the training regime, Chronologer is designed to assume a fixed peptide-level FDR and learns to discard a fraction of training data that disagrees with the consensus. Chronologer is trained using Chronologer-DB, a vast 2.2 million peptide training set generated from eleven community datasets with ten different types of PTMs. These datasets have been harmonized to a common scale, enabling a single model that predicts all PTM types. We find that the scale of Chronologer-DB dramatically improves the accuracy of Chronologer to produce less than two-thirds the error rate of other deep-learning predictors. Using a new generalized approach for transfer learning, we demonstrate that Chronologer can robustly learn the effects of rare and chemically unique modifications present on as few as 100 peptide observations while retaining the ability to predict previously learned peptide classes.

## Results and Discussion

### Chronologer architecture and evaluation

Chronologer is a new deep-learning architecture to predict C18 RP-HPLC RTs from peptide sequence alone. Inspired by recent innovations in protein secondary structure prediction^24^ and functional classification,^25^ Chronologer learns to encode each amino acid type as an array of 64 numbers that, when tiled along a sequence dimension, creates a 2-dimensional matrix representation for each peptide. A series of linear transforms are performed on these matrices in a sliding window fashion (convolutions) to incorporate sequence information, with dilations of the window size (kernel) allowing gradual incorporation of longer distance content. The architecture includes skip functions to allow passive information transfer and preserve the learning gradient as additional numerical operations are performed (**Fig 1A**).

**Figure 1:**
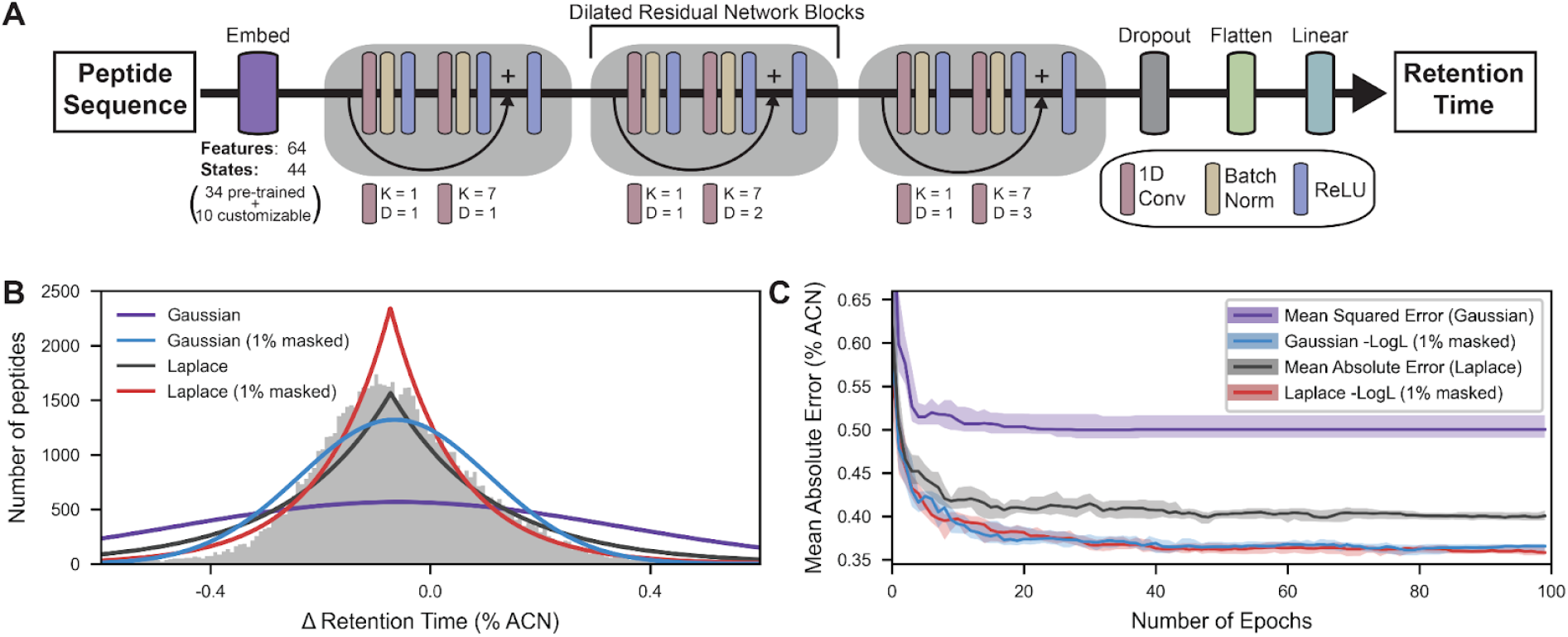
Chronologer architecture and training. (**A**) Chronologer uses a residual convolutional neural network (RCNN) architecture where peptide sequences are numerically encoded using a learned embedding layer (64 features), processed through a series of residual network blocks, and C18 RP-HPLC RT decoded using a fully connected linear layer. Each residual block contains a feature generation step – a 1D convolution of kernel (K) size 1 – followed by a bottleneck convolution (K=7) with increasing dilation rates (D) to gradually incorporate longer distance sequence information. Residual skip functions (arrows) improve information flow through the model and stabilize the gradient for model training. A single dropout layer is included for model regularization. (**B**) Comparison of empirical RT errors (histogram) for a single HeLa lysate analyzed on different LC systems with optimized fits for both Gaussian (mean squared error) and Laplace (mean absolute error) distributions with or without 1% outlier masking. (**C**) Using the Brurderer et al. (2017) dataset, Chronologer was trained using both Gaussian- and Laplace-based loss functions with or without 1% outlier masking that functions as a FDR-filter; irrespective of base distribution, masking substantially improves the performance on holdout data during training.

The model parameters of Chronologer are trained by minimizing the output of a new loss function that compares real and predicted RTs. We developed this loss function based on the empirical properties observed in comparisons of RTs for a single preparation of tryptic peptides from HeLa cell lysate analyzed on multiple columns, LC systems, and mass spectrometers. Most machine learning methods assume errors are Gaussian distributed, where the distribution spread is proportional to the mean-squared error. For training purposes, precisely fitting the center of these distributions matters less than fitting the extreme values for modeling errors. We found that for these regions, differences in peptide RTs are better described by a Laplace distribution (**Fig 1B**) whose spread is proportional to the mean absolute error (MAE).

Additionally, Chronologer training includes a mechanism to account for errors in the underlying training data. While data for machine learning is typically assumed to be error-free, most peptide datasets are created with an implicit 1% false discovery rate (FDR) and we expect a similar percentage of erroneous observations in our training data. These incorrectly assigned peptide RTs will usually result in errors that fall outside of the expected Laplace distribution and such outliers have an outsized influence on model parameters during training (e.g. if we have a correct observation with 30 sec RT error and an erroneous observation with a 10 min RT error, the model optimizer will give 20-fold more weight to the erroneous observation). We found that a simple and effective solution to minimize the contribution of false discoveries was to dynamically mask any data that falls outside of the inferred 99% confidence interval during training (**Fig 1C**). The benefits of FDR-masking are even more pronounced when predicted RTs are assumed to be Gaussian distributed, where outliers have a larger impact on the gradient due to squaring of the residual terms in place of absolute residuals in the Laplace distribution.

We assessed the performance of the Chronologer architecture and loss function by training independent models with three large datasets (composed of ∼320-450K unique peptides each) that were used to train three different contemporary RT predictors: Prosit,^17,26^ DeepLC,^19^ and SSRCalc.^15,27^ Chronologer outperformed all three tools by 23-39% when trained on the same source data (**Fig S1**), which supports that our new architecture and loss function are effective strategies for C18 RP-HPLC RT prediction.

### Aggregating and harmonizing community-wide datasets

While evaluating the performance of Chronologer, we were surprised to find that the three large RT datasets used for training were largely non-redundant, and collectively contained nearly 1 million unique peptide RT measurements (**Fig S2**). The performance of deep learning models is highly dependent on the breadth and scale of training data, and we sought to combine these and other large proteomics datasets (11 datasets with 10 modification types, see **Table 1**) into a single database for Chronologer training. As each peptide dataset was collected using a different HPLC system, C18 resin, and chromatography gradient, individual RTs between datasets are not directly comparable and needed to be aligned into a common RT space for comparison.

**Table 1:** Datasets compiled in Chronologer-DB.

| Reference | Description | # unique peptides | # Prosit-predictable peptides | Name in Fig 2 |
| --- | --- | --- | --- | --- |
| Bruderer et al.<sup>28</sup> | Deep HeLa library | 321,640 | 309,614 | HeLa |
| Schmidt et al.<sup>23</sup> | ProteomeTools synthetic library of human-like peptides, mostly tryptic | 458,212 | 457,006 | PT-Tryptic |
| Mizero et al.<sup>27</sup> | Training data from the SSRCalc database | 400,170 | 392,716 | % ACN |
| Wilhelm et al.<sup>26</sup> | ProteomeTools synthetic library of HLA peptides | 719,888 | 718,104 | PT-HLA |
| Miller et al.<sup>29</sup> | Jurkat cell lysate treated with six different proteases | 104,300 | 95,312 | Multi-protease |
| Cappelletti et al.<sup>30</sup> | Limited tryptic proteolysis of *Escherichia coli* and *Saccharomyces cerevisiae* lysates | 46,691 | 45,570 | Long peptides |
| Lawrence et al.<sup>31</sup> | Phosphopeptides enriched from tryptic proteolysis of MCF7 cell lysate | 212,679 | 270 | Phospho |
| Hanson et al.<sup>32</sup> | Enrichment of ubiquitylated tryptic peptides (using anti-ε-KGG) from HEK293 and U2OS cell lysates | 145,491 | 52,084 | Ubiquitination |
| Baeza et al.<sup>33</sup> | Peptides from tryptic proteolysis of *in vitro* acetylated MCF7 cell lysate | 15,463 | 5,644 | Acetylation |
| Robinson et al.<sup>34</sup> | Synthetic and natural mouse liver tryptic peptides with mono/di/(tri)-methylated Lys and Arg | 38,450 | 35,929 | Methylation |
| Zecha et al.<sup>35</sup> | Sub-stoichiometric TMT labeled tryptic peptides from HEK293, Jurkat, and mouse liver lysates | 172,707 | 2,644 | TMT |

Peptide datasets are generally aligned using standards common to both datasets, which may be either naturally shared by the samples^12,31^ or added exogenously as synthetic spike-in standards^11^ (**Fig 2A**). Linearity is assumed between retention spaces when only a few shared peptide standards are available (e.g. the 11 synthetic iRT peptides in Escher et al.^11^), but this assumption is rarely true due to differences in elution gradients and HPLC systems. Nonlinear regression is possible when 100s-1000s of standards are available. Without synthetic standards, many datasets share no peptides; for example, when samples are from different species, enriched for post-translational modifications (PTMs), or proteolyzed with alternative proteases (**Fig S3**).

**Figure 2:**
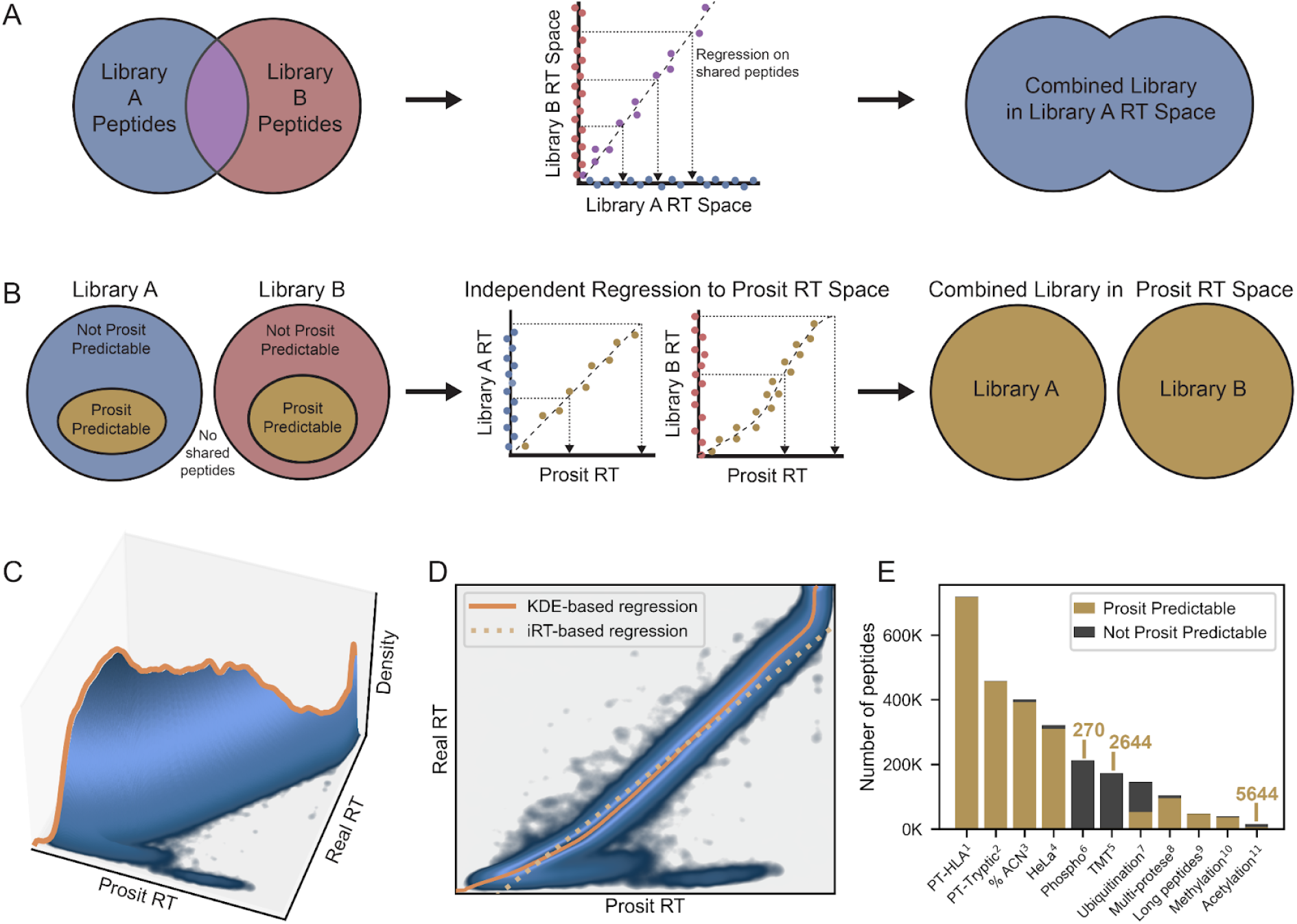
Generation of the Chronologer Database. (**A**) Traditional alignment of peptide RT libraries requires using shared peptides present in both libraries (such as spiked iRT standard peptides) to provide a mapping between RT spaces; linear regression is standard when the number of shared peptides is limited (e.g. 11 iRT standards). (**B**) We propose a new strategy, *in silico*-based alignment, that leverages existing deep learning tools to highly diverse libraries that do not necessarily contain any shared peptides. Nearly all peptide libraries possess hundreds to thousands of peptides that can be predicted by Prosit (7-30 amino acids with fixed C carbamylation and variable M oxidation) which provide references for non-linear mapping into a common Prosit RT space. (**C-D**) KDE-based regression into Prosit RT space is a nonparametric approach that is highly robust to outliers and outperforms iRT-based linear regression, as demonstrated when aligning Prosit predicted RTs to empirical observations from Bruderer et al.^28^ (**E**) Eleven large datasets with diverse peptide chemistries were aggregated using *in silico*-based alignment to produce the 2.2M peptide Chronologer database (annotations are provided for datasets with the fewest Prosit predictable peptides).

A key aspect of RT alignment is that it does not alter the elution order of the peptides and only serves to rescale a given dataset onto a different reference dimension. While iRT units or an experimental library are common choices as the “reference,” the exact choice of the reference dimension is fairly arbitrary. Therefore, we propose a modified alignment strategy that does not rely on shared peptides but rather leverages the availability of peptides whose RTs can be predicted *in silico*. Since PTM-enriched datasets also contain hundreds or thousands of unmodified peptide signals, this approach allows us to align practically any dataset into a common reference RT space by non-linear methods (**Fig 2B**). To do this, we used the deep learning tool Prosit which can predict unitless relative RTs for peptides that are 7-30 residues in length with fixed cysteine carbamidomethylation and variable methionine oxidation. We have found that peptides with these characteristics are common, even in datasets that are highly enriched for select PTMs.

The kernel density estimation (KDE) based alignment algorithm^6^ in EncyclopeDIA can trace a monotonic function through RT space following the path of highest peptide density (**Fig 2C-D**). This KDE algorithm has been demonstrated robust to outliers from false peptide identifications as well as correlated errors in Prosit predictions.^36^ All 11 large peptide datasets contained enough Prosit predictable peptides for KDE-based alignment into Prosit RT space (**Fig 2E**), enabling the construction of the new 2.2 million peptide Chronologer database (Chronologer-DB). For context, the Chronologer-DB is a RT library of comparable scale to MassIVE-KB for mass spectra (∼2.5 million peptides).^37^ The Chronologer-DB has been aligned into % acetonitrile space^38,39^ using the SSRCalc dataset, which allows Chronologer to generate predictions in biochemically meaningful units. When trained using this new massive database, Chronologer outperforms the four evaluated tools by 36-65% with more consistent variation across a standard 90-min gradient (**Fig 3A-B**). We also observe that Chronologer predictions increased the number of detections when library searching with Skyline^40,41^ and mProphet^42^ relative to other RT prediction models and performed most closely to empirical RTs collected on the same LC system with the same column and buffers (**Fig 3C**).

**Figure 3:**
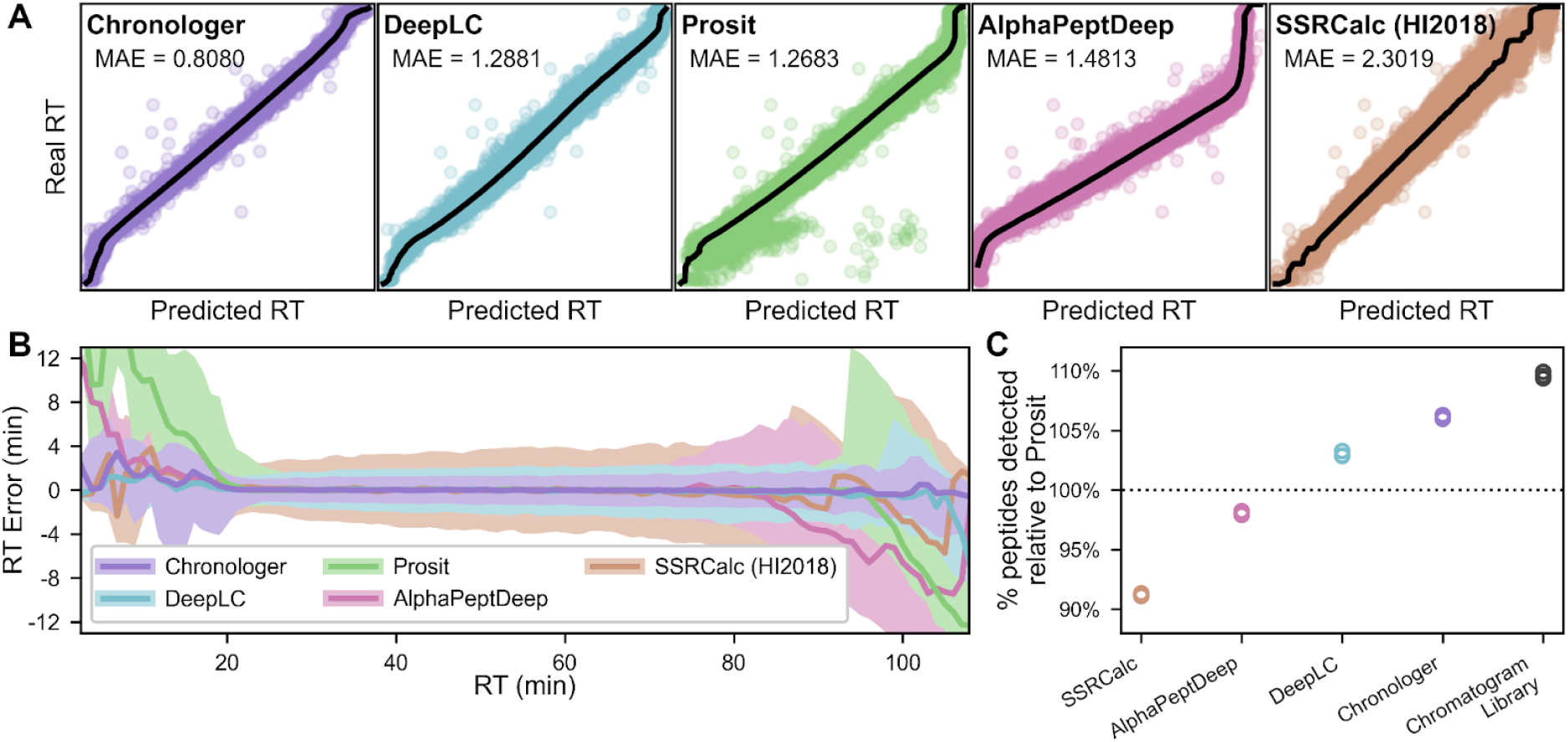
Chronologer predictions are more accurate than other tools across gradients. (**A**) HeLa peptides from Searle et al.^6^ were aligned using the KDE algorithm to predictions by Chronologer, DeepLC, Prosit, AlphaPeptDeep, and SSRCalc, with Chronologer having the lowest mean absolute error (highest accuracy) relative to the regression line (black). (**B**) The distribution of RT errors across the gradient for each prediction method averaged over 1 min sliding windows with the median plotted as a solid line and shading denoting the middle 90% percentile. Chronologer predictions are generally more consistent across the gradient compared to other tools. (**C**) The relative increase or decrease between prediction tools (relative to Prosit) of the number of detected HeLa peptides analyzed using Skyline/mProphet considering the same spectrum library but with different predicted RTs. The chromatogram library measurements represent gold-standard RTs empirically measured on the same column.

As the Chronologer database includes greater peptide diversity than any single existing dataset, we found that performance gains with Chronologer are consistent across different proteolytic digests, variations in column temperature, and even outperform specialized models designed for specific PTMs, such as DeepPhospho (**Fig 4**). Compared to the other RT prediction tools, DeepLC is unique as it includes a chemical structure model to predict RTs for “as-yet-unseen” modifications that are not included in its training data, such as phosphopeptides. However, as there are no phosphates in any of the DeepLC training data, unsurprisingly we find that it performs substantially worse for this PTM compared to models trained on real observations (Chronologer, DeepPhospho, AlphaPeptDeep).

**Figure 4:**
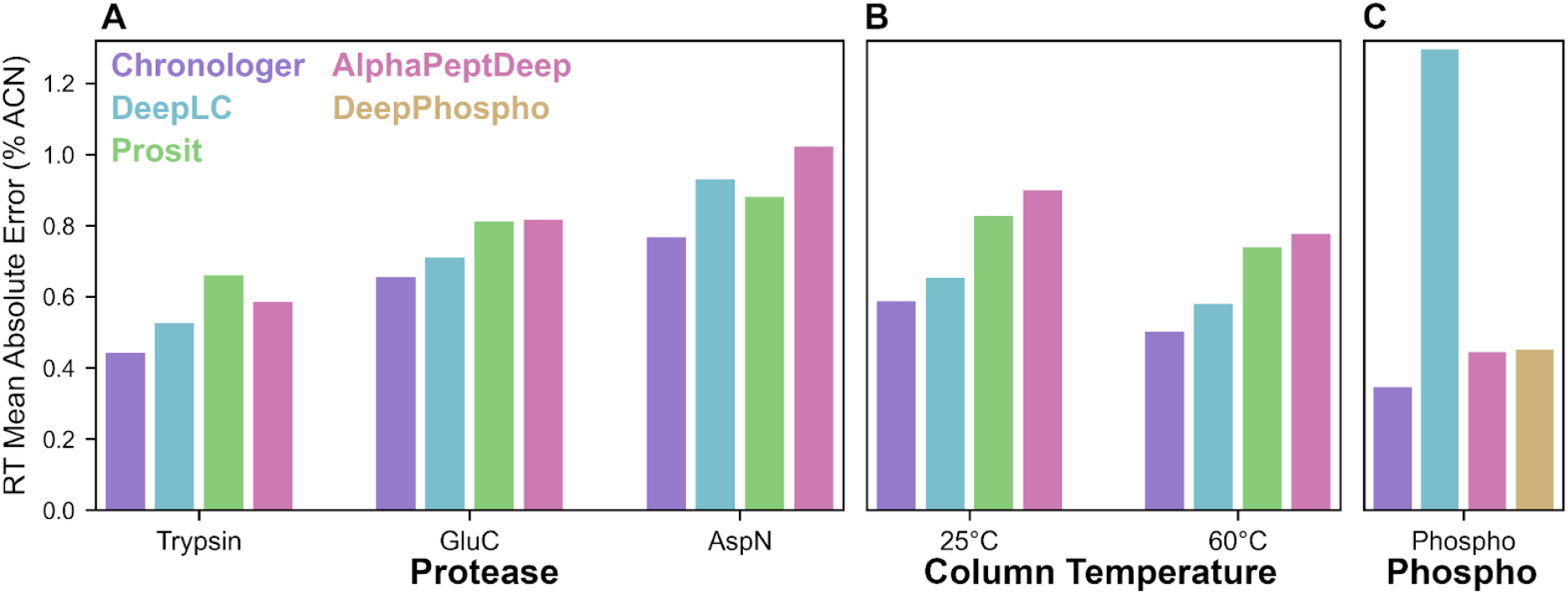
Chronologer performs well across diverse sample types. When compared to other tools, Chronologer consistently produced the most accurate RT predictions for (**A**) peptides produced from different proteases digests.^43^ (**B**) separations at different column temperatures,^44^ and (**C**) phosphopeptides.^45^

### Extending Chronologer to predict retention times of rare post-translational modifications

While the current Chronologer database is extensive in scale, it does not contain an exhaustive sampling of all peptide modification types. Neural networks are often re-trained with additional data to improve their performance for specific applications, referred to as “transfer learning”.^20,46^ As Chronologer has already been trained across a wide breadth of peptide chemistries via the alignment of independent libraries, we chose to evaluate how efficiently Chronologer could learn new peptide modification types when training was augmented with additional data.

As a general test of the approach, we first performed a series of experiments where Chronologer was trained on a large library of peptides (400K) which contained only 19 of the 20 common amino acids, and then re-trained with an augmented library that included a variable number of peptides containing the 20th amino acid. For most residue types, as few as 100-1000 new peptides containing that residue were sufficient for Chronologer to generate accurate predictions of that amino acid (**Fig 5**). It was notable that proline, which can undergo cis-trans isomerization, was the most difficult amino acid for Chronologer to learn and required 10K-100K peptides to achieve similar performance. Importantly, re-training with data containing an additional amino acid did not alter the quality of predictions on peptides absent of that residue type. These synthetic experiments suggested that Chronologer can efficiently learn new peptide chemistries with limited observations.

**Figure 5:**
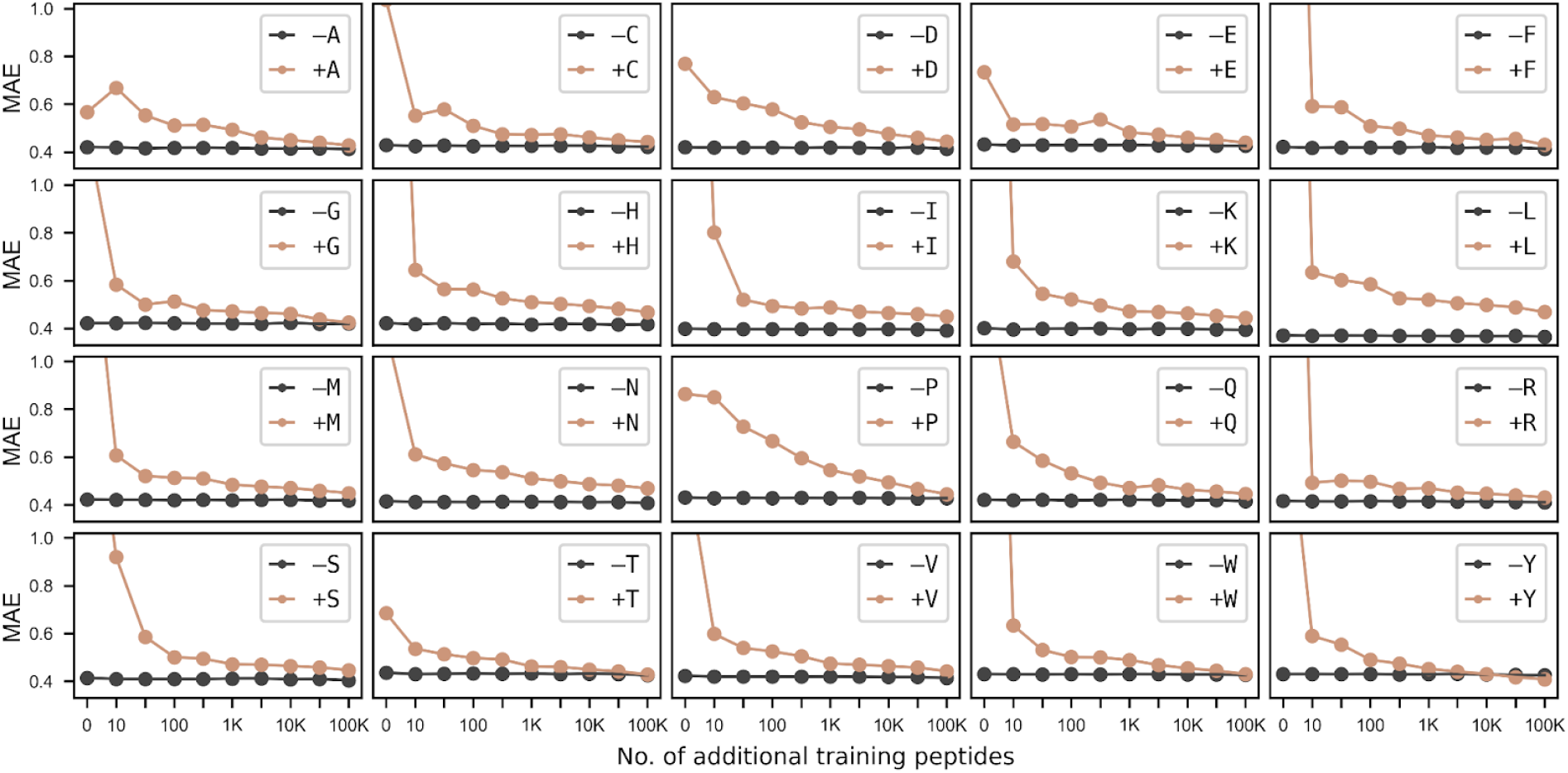
Chronologer can learn new amino acid types with limited training data. A synthetic experiment where Chronologer was pre-trained using peptides that contained only 19 of the 20 amino acids, followed by training that was augmented with a limited number of peptides containing the 20th amino acid. With few exceptions (notably proline), Chronologer could learn to accurately predict the properties of the 20th amino acid with as few as 10-100 examples.

As a more realistic demonstration, we evaluated the efficiency of re-training Chronologer to predict peptides with S/T residues modified with OGlcNAc, a small O-glycan that has been functionally associated with the regulation of apoptosis,^47^ nutrient sensing,^48^ oxidative stress,^49^ and proteostasis.^50^ The labile nature and low antigenicity of OGlcNAc make it a difficult modification to study through targeted enrichment, and as such, there are relatively few site-localized observations of OGlcNAc-modified peptides. Additionally, since Chronologer has never been exposed to glycans of any kind, studying this moiety demonstrates how the algorithm deals with truly new PTMs. Using recently published data by Burt et al.,^51^ Chronologer was first trained with the full 2.2M peptide database and then re-trained with augmentation of OGlcNac peptides found in mouse synaptosomes (10-1473 OGlcNAc peptides); prediction performance was then evaluated using independently acquired mouse embryonic stem cell data (507 OGlcNAc peptides). With only ∼100 peptides added to the training data, Chronologer learned how to accurately predict RTs (MAE = 0.67% ACN) for OGlcNAc peptides (**Fig 6A**), which is remarkable given that no other glycans are present in the database. Using the entire 1473 OGlcNAc peptides with iterative training, Chronologer achieved a MAE of 0.48% ACN, which was 3.6x lower than comparable predictions by DeepLC (MAE = 1.73% ACN).

**Figure 6:**
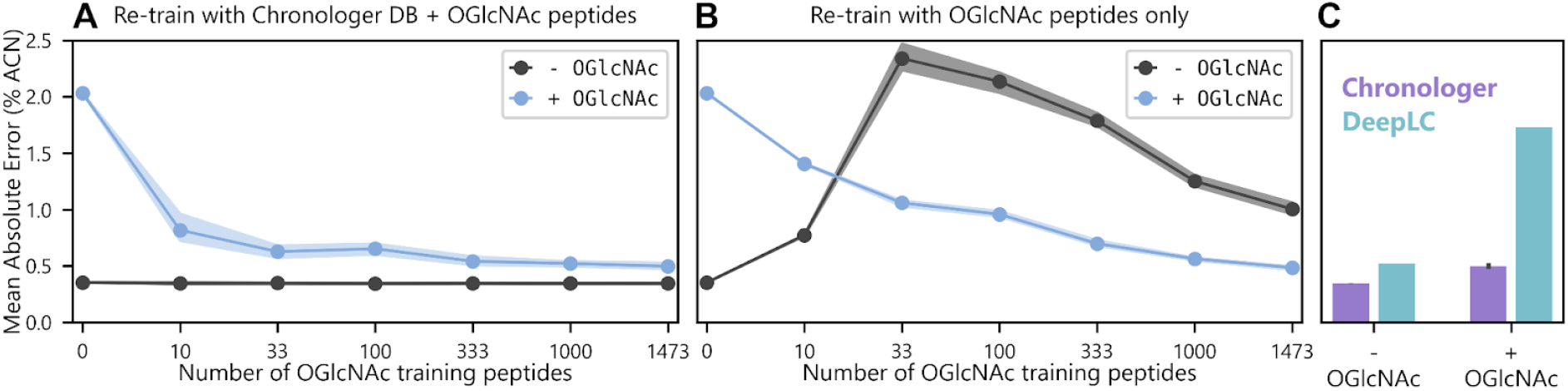
Chronologer can learn chemically distinct modifications like glycans with limited training data. (**A**) Performance of evaluation peptides from mouse embryonic stem cell data with or without OGlcNAc modifications using Chronologer after retraining using the Chronologer-DB augmented with a limited number of OGlcNAc peptides (30 replicates, with points denoting the median and shading for the middle 80%ile) (**B**) Chronologer performs worse on both -/+ OGlcNAc peptides if re-training is performed using only the new limited OGlcNAc data absent the Chronologer-DB and highlights the utility of data harmonization compared to overfitting to limited observations. (**C**) When trained using a relatively small number of OGlcNAc-containing peptides (n=1473), Chronologer outperforms the DeepLC predictions that use a chemical structural model with no OGlcNAc training data.

Interestingly, if Chronologer is re-trained with only the OGlcNAc peptides absent from the original 2.2M peptide database, it both quickly forgets how to accurately predict unmodified peptides and also does not as efficiently learn OGlcNAc chemistry (**Fig 6B**). From this, we conclude that the model is quick to overfit to new data when lacking more diverse chemical constraints provided by the massive Chronologer-DB. OGlcNAc prediction quality was similar between our augmentation training method and complete retraining of Chronologer with both the Chronologer-DB and OGlcNAc peptides (**Fig S4**). Interestingly, despite the fact that OGlcNAc contains atoms and chemical functional groups that match those found in the chemical structure-based model of DeepLC, it significantly underperformed relative to OGlcNAc-augmented Chronologer (**Fig 6C**), even when training included as few as 10 OGlcNAc peptides. This augmentation training method represents a different approach to transfer learning that requires very little new data since the model is added to, rather than fully re-trained.

## Conclusion

In summary, Chronologer is a new deep learning package for accurate prediction of C18 RP-HPLC RTs from peptide sequences with support for 10 modification types and easy upgradability for new modifications requiring only limited data. Chronologer makes predictions using a simple convolutional network that is built around a framework of data harmonization with empirically-informed priors for erroneous data rates and expected chromatographic errors. Like all regression methods, the generalizability of deep learning models is highly dependent on the quality of training data, and the size of Chronologer-DB helps Chronologer robustly generalize across a wide range of peptides by leveraging learning from many experimentally distinct datasets. The simple architecture allows rapid execution and augmented training of Chronologer using commodity hardware such as laptops that lack a GPU. The Chronologer dataset and code base are freely available and fully open source.

Machine learning methods generally consider all training data to be correct, which can result in overtraining around dataset artifacts. Using our likelihood-based loss function with a tunable FDR, Chronologer naturally learns to filter likely faulty observations that would otherwise influence model parameters and bias the prediction performance. Finally, our augmentation training method repurposes transfer learning to improve learning globally, rather than on a new, limited dataset. We believe these new approaches to training with implicit dataset FDR have generalizable extensions in any machine learning situation where training data cannot be hand-curated.

## Acknowledgements

This research was supported by NIH R01GM133981 to DLS and BCS, and NIH R00HD090201 to DBW.

## Author Contributions

DBW and BCS conceived the study. DBW, AES, VS, ALR, and DY performed the experiments. DLS, OVK, and BCS supervised the work. DBW and BCS wrote the manuscript; all authors edited and approved the work.

## Competing interests

The authors declare the following competing interests: BCS is a founder and shareholder in Proteome Software, which operates in the field of proteomics. DLS has a consulting agreement with Maze Therapeutics.

## Data/code availability

The code for chronologer is publicly available at https://github.com/searlelab/chronologer under the Apache 2.0 open source license.

## Materials and Methods

### Kernel density estimation (KDE)-based retention time alignment method

Unless stated otherwise, alignment between retention libraries was performed using a Python implementation of the KDE-based algorithm within EncyclopeDIA.^6^ Nearly all peptide libraries contain some number of erroneous detections (usually at ∼1% FDR) that will skew regression lines fit by minimization approaches whose loss functions are sensitive to the magnitude of outliers (e.g. least squares). By contrast, the EncyclopeDIA alignment algorithm is a non-parametric method that produces a monotonic function to interconvert between two RT spaces based on their modal point densities and is highly robust to outliers. Briefly, a 2-dimensional kernel density map of the two RT spaces is generated by summing kernel estimates for each peptide common to both libraries. Kernel estimates are 2-dimensional Gaussian-like curves centered on the observed RTs in each library with a standard deviation based on Silverman’s rule:

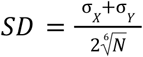

Where N is the number of shared peptides and σ_X_, σ_Y_ are the standard deviations of each RT library. For computational efficiency, the kernel density map is defined as a two-dimensional array with 3,000 points along each RT dimension and kernel estimates use a cosine-approximate Gaussian function. A modal line is traced through the two RT spaces by monotonically following the path of maximum density; specifically, the 2-dimensional index with the global maximum density is identified, and iterative steps are performed in both the positive and negative directions to identify a path that proceeds to the point of next highest density. A spline function with smooth linear interpolation is fit to the identified modal points using the Scipy interp1d function.^52^

### Chronologer database construction by *in silico* retention time alignment

The Chronologer database was constructed through the alignment of eleven RT datasets that together represent ∼2.2 million unique peptide sequences with ten different types of modifications. These libraries were selected to provide adequate training data across a range of chemical conditions to improve the robustness of Chronologer conditions. Bruderer et al.^28^ (321,640 peptides), Schmidt et al.^23^ (458,212 peptides), and a large dataset developed training SSRCalc^27^ (400,170 peptides) are primarily composed of unmodified tryptic peptides. The synthetic HLA peptide library of Wilhelm et al.^26^ (719,888 peptides) and the multiple protease library of Miller et al.^29^ (104,300 peptides) substantially increase the number of non-tryptic peptides. A limited proteolysis library by Cappelletti et al.^30^ (46,691 peptides) includes many longer peptides up to 50 amino acids. Five modification-specific libraries were also included: Lawrence et al.^31^ (212,679 peptides) for phosphorylation, Hanson et al.^32^ (145,491 peptides) for ubiquitylation, Baeza et al.^33^ (15,463 peptides) for acetylation, Robinson et al.^34^ (38,450 peptides) for methylation, and Zecha et al.^35^ (172,707 peptides) for TMT labeling. Each library was mapped into Prosit RT space through KDE-based alignment using all available Prosit predictable peptides (i.e. peptides between 7 and 30 amino acids with fixed cysteine carbamidomethylation and variable methionine oxidation). For 5 of the datasets (Wilhelm, Miller, Hansen, Baeza, and Zecha), peptide-to-RT tables were not readily available for peptide RTs and/or Prosit-predictable peptides were not directly reported in the manuscripts. For these datasets, raw files were re-searched using MSFragger v3.2 with the appropriate sequence database (synthetic HLA peptides or Uniprot reference human/mouse proteomes downloaded 2022-3-02) and similar search parameters to the original published settings (**Table S1**).

Non-redundant spectral libraries were then prepared using BlibBuild and BlibFilter within Bibliospec 2.0^53^ and converted to DLIB format with Encyclopedia v.1.12.31^6^ for subsequent data processing. All datasets were then mapped from the arbitrary Prosit RT space to the more physiochemically relevant hydrophobic index (HI): the % acetonitrile (in 0.1% formic acid) at which a peptide elutes by C18 RP-HPLC.^39^ The SSRCalc dataset was experimentally calibrated in HI space, and a reciprocal function was computed by KDE-based alignment to map the Prosit RT-aligned Chronologer database into HI space. Consequently, models trained using the Chronologer database provide a prediction of peptide elution in chemical units of fractional mobile phase.

### Chronologer architecture, model training, and evaluation

Inspired by the work of Bileschi et al.,^25^ the Chronologer deep learning model predicts peptide RTs using a residual convolutional neural network (RCNN) that was implemented using Pytorch.^54^ Briefly, the Chronologer model consists of an embedding layer with 64 learned amino acid features, a series of three residual dilated convolution blocks for encoding sequence information, and a single linear decoder layer to predict RT. The model is regularized during training by random dropout (10%) of encoded parameters prior to decoding. Each residual block consists of a feature generation step (1D convolution with kernel size 1, normalization, and activation), a bottleneck step (same series with an expansion of the kernel size to 7), a skip function to enable passive information transfer, and a final activation step. Chronologer parameters are trained using gradient-based optimization with Adam^55^ and learning rate modulation based on Smith et al.^56^ where the learning rate is fixed (0.001) and the batch size is modulated (64×2^*e*/30^ for each epoch *e*, rounded to the nearest multiple of 8). Twenty percent of data was withheld for model evaluation. Chronologer was evaluated against competing tools (DeepLC, Prosit, SSRCalc, DeepPhospho) using tryptic HeLa peptides by Searle et al.,^6^ a multi-protease panel by Richard et al.,^43^ tryptic peptides separated at different column temperatures by Li et al.,^44^ and phosphopeptides from HeLa lysate by Searle et al..^45^ Prosit predictions were made using the 2019 iRT model through the available ProteomicsDB web server; DeepLC predictions were performed locally with v.1.1.2 and the pre-trained model based on Bruderer et al.;^28^ DeepPhospho predictions were performed locally using the 20211212 build with the included pre-trained RT model.

### Chronologer loss function optimization

Maximum likelihood parameter estimates of Gaussian and Laplace distributions (with or without masking of outliers outside the inferred 99% confidence interval) were computed using Scipy.^52^ The same distributions (Gaussian and Laplace, with and without 1% FDR masking) were evaluated as loss functions during Chronologer training using data from Bruderer et al. (2017). Gaussian and Laplace distributions were optimized as mean squared error and mean absolute error, respectively, with FDR-masked variants using the negative log-likelihood as a loss function. Full Chronologer training was performed using a Laplace-based negative log-likelihood function with 1% FDR masking and independent error rates for each constituent dataset of the Chronologer database.

### Evaluation of Chronologer re-training

We evaluated the ability of Chronologer to learn new amino acid chemistries through two experiments. First, in a “leave one out” design, we performed a synthetic experiment where Chronologer was trained on 400K unmodified peptides randomly sampled from the Chronologer database that contained only 19 of the 20 common amino acids (using the standard Chronologer training regime), followed by re-training with the data augmented to include 10 to 100K peptides that also include the 20th amino acid (same training protocol except the initial batch size was doubled to halve the learning rate). The performance of these models was evaluated on an independent set of 44K and 11K peptides that excluded or included the 20th amino acid, respectively, Second, we evaluated how efficiently a fully trained Chronologer model could learn to OGlcNAc modifications on S and T residues using data from Burt et al.^51^ that included two datasets with high confidence, site-localized OGlcNAc peptides: a mouse embryonic stem cell library and mouse synaptosome library with 507 and 1569 OGlcNAc peptides and OGlcNAc peptides, respectively. Shared peptides were filtered from the synaptosome library – leaving 1473 peptides – which were used for model training and the embryonic stem cell dataset reserved for model evaluation. We evaluated three different training regimes where Chronologer was presented with a limited number of OGlcNAc peptides (10-1473). First, following routine pre-training with ChronlogerDB, the database was augmented with OGlcNAc peptides and training continued with twice the initial batch size (same as the leave-one-residue-out experiment). Second, after ChronologerDB pre-training, the model was presented with only OGlcNAc peptides (no ChronologerDB). And third, models were fully trained with ChronologerDB and OGlcNAc peptides from random starting parameters and no pre-training.

### Library searching using predicted retention times

Validation of the Hela replicate dataset^36^ was performed according to methods previously used to compare DDA and DIA datasets with Skyline^40,41^ version 22.2. Separate BLIB and iRTDB libraries from the same 98,519 peptides with identical MS2 fragmentation patterns were generated using retention times from SSRCalc, AlphaPeptDeep, Prosit, DeepLC, or Chronologer. In all cases, 69 high abundance peptides were selected as iRT anchors for retention time alignment. For all libraries, Skyline was configured to import precursor isolation windows from the result files from 388 to 1000 m/z, where peptides were allowed to contain charges from +2 to +4. Chromatograms for ladder fragment ions (excluding y1, b1, y2, and b2) from 300 to 2000 m/z were extracted, where up to 9 transitions (6 MS2 and 3 MS1) were extracted. MS1 and MS2 extractions were configured to find signal within a 10 ppm mass error from centroided, demultiplexed spectra. A mProphet^42^ model was trained using second-best peaks, and applied excluding run-to-run alignment.

