## Supplementary Materials for "Deep learning from harmonized peptide libraries enables retention time prediction of diverse post translational modifications"

### Supplementary Figures:

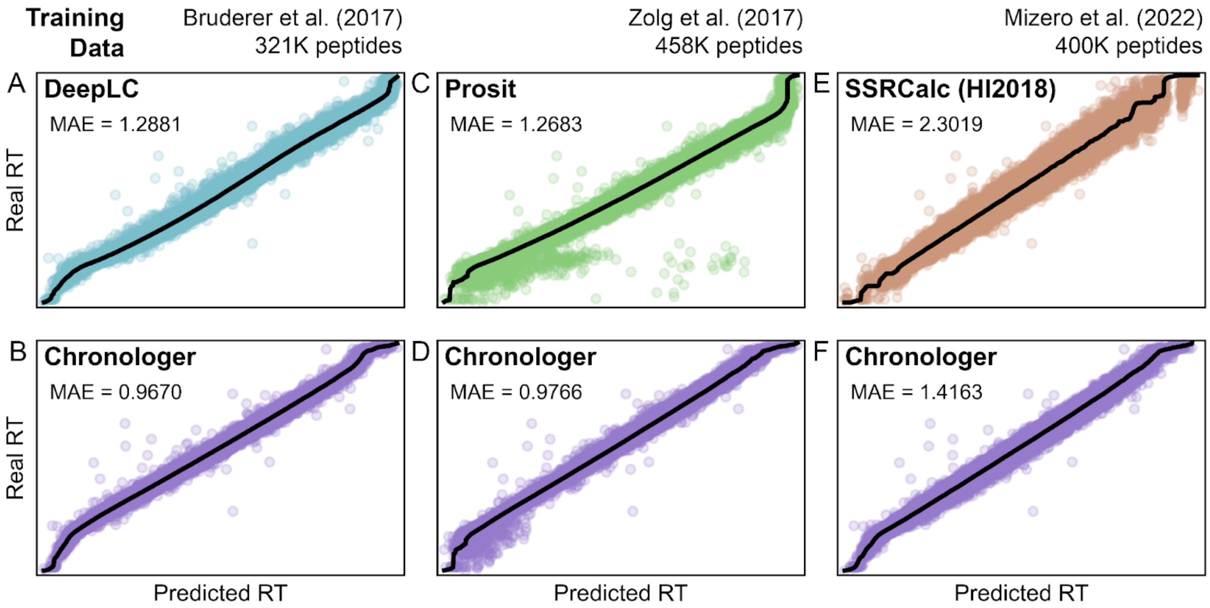


**Figure S1: Chronologer performance on different datasets**. The performance of the Chronologer architecture was compared to three popular RT prediction models using the same underlying training data: (**A-B**) DeepLC with Bruderer et al. (2017), (**C-D**) Prosit with Zolg et al. (2017), and (**E-F**) SSRCalc with Mizero et al. (2022). When trained using each of the underlying datasets, the Chronologer architecture performs substantially better than the previous tools.

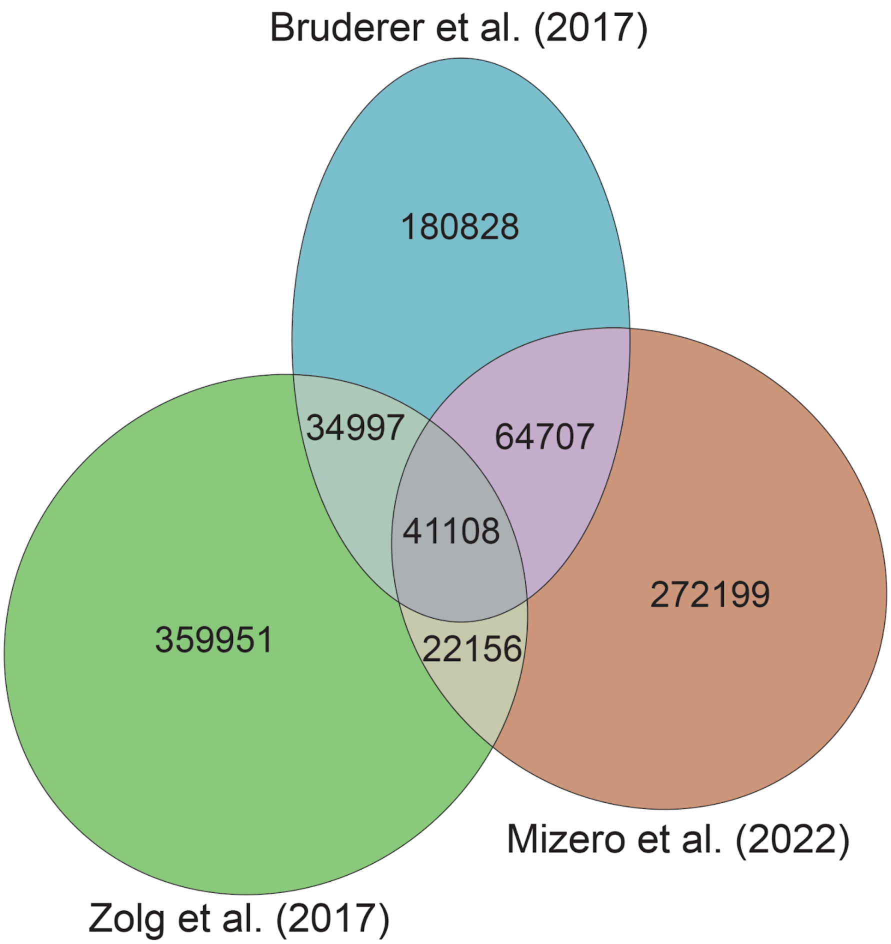


**Figure S2: Shared peptides between large tryptic peptide libraries**. The large libraries used to train DeepLC, Prosit, and SSRCalc are largely non-overlapping in the peptides present.

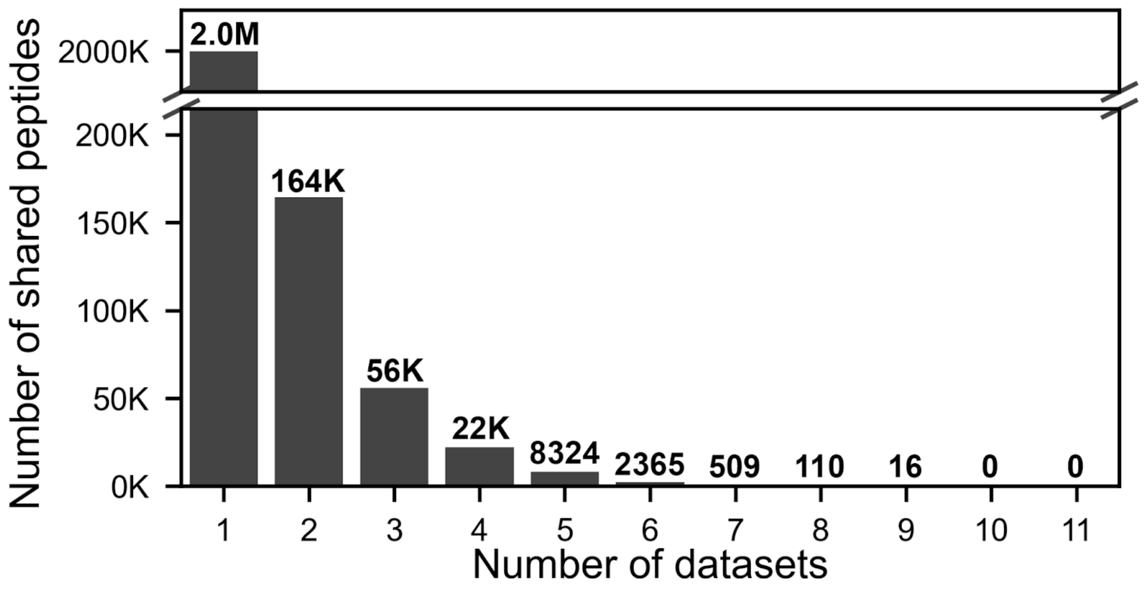


**Figure S3: Shared peptides between datasets in the Chronologer-DB**. Most peptides in the Chronologer-DB are only observed in only a single dataset and no single peptide is observed in all constituent datasets.

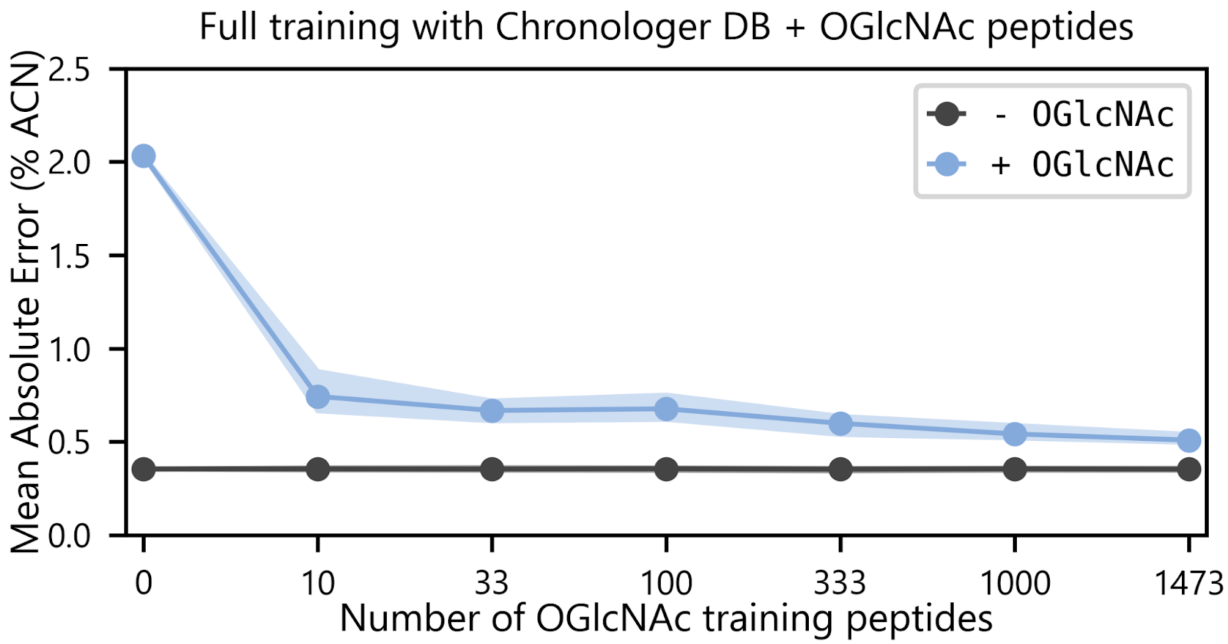


**Figure S4: OGlcNAc training with random starting parameters**. Full training of Chronologer where both the Chronologer-DB and OGlcNAc peptides were present shows no difference in final model performance compared to augmented training where OGlcNAc peptides are added after initial model training with the Chronologer-DB.

**Table S1: Search engine settings used to analyze datasets in Chronologer-DB**

| Reference | MSFragger parameters |
|---|---|
| Wilhelm et al.[25] | Precursor mass tolerance = 50 ppm<br>Fragment mass tolerance = 20 ppm<br>Protease = Nonspecific<br>Allowed miscleavages = 0<br>Fixed modifications: C +57.021464<br>Variable modifications: M +15.994900<br>nQnC -17.026500<br>nE -18.010600<br>Max variable modifications = 3 |
| Miller et al.[43] | Precursor mass tolerance = 50 ppm<br>Fragment mass tolerance = 20 ppm<br>Protease = Trypsin (KR/P), LysC (K/P), ArgC (R/P), Chymotrypsin (FLWY/P), AspN (D), GluC (DE/P), matched to the dataset<br>Allowed miscleavages = 2<br>Fixed modifications: C +57.021464<br>Variable modifications: M +15.994900<br>N-term +42.010565<br>nQnC -17.026500<br>nE -18.010600<br>Max variable modifications = 3 |
| Hanson et al.[45] | Precursor mass tolerance = 50 ppm<br>Fragment mass tolerance = 20 ppm<br>Protease = Trypsin (KR/P)<br>Allowed miscleavages = 3<br>Fixed modifications: C +57.021464<br>Variable modifications: M +15.994900<br>N-term +42.010565<br>nQnC -17.026500<br>nE -18.010600<br>K +114.042927<br>Max variable modifications = 3 |
| Baeza et al.[46] | Precursor mass tolerance = 50 ppm<br>Fragment mass tolerance = 20 ppm<br>Protease = ArgC (R/P) + GluC (E/P)<br>Allowed miscleavages = 2<br>Fixed modifications: C +57.021464<br>K +42.015650<br>Variable modifications: M +15.994900<br>N-term +42.010565<br>nQnC -17.026500<br>nE -18.010600<br>Max variable modifications = 3 |

| | |
|---|---|
| Zecha et al.[48] | Precursor mass tolerance = 50 ppm<br>Fragment mass tolerance = 20 ppm<br>Protease = Trypsin (KR/P)<br>Allowed miscleavages = 2<br>Fixed modifications: C +57.021464<br>Variable modifications: M +15.994900<br>N-term +42.010565<br>nQnC -17.026500<br>nE -18.010600<br>K +224.152478 (TMT0) or K +229.162932 (TMT10)<br>Max variable modifications = 3 |
